# Tightly constrained genome reduction and relaxation of purifying selection during secondary plastid endosymbiosis

**DOI:** 10.1101/2021.05.27.446077

**Authors:** Kavitha Uthanumallian, Cintia Iha, Sonja I. Repetti, Cheong Xin Chan, Debashish Bhattacharya, Sebastian Duchene, Heroen Verbruggen

## Abstract

Endosymbiosis, the establishment of a former free-living prokaryotic or eukaryotic cell as an organelle inside a host cell, can dramatically alter the genomic architecture of the endosymbiont. Plastids, the light harvesting organelles of photosynthetic eukaryotes, are excellent models to study this phenomenon because plastid origin has occurred multiple times in evolution. Here, we investigate the genomic signature of molecular processes acting through secondary plastid endosymbiosis – the origination of a new plastid from a free-living eukaryotic alga. We used phylogenetic comparative methods to study gene loss and changes in selective regimes on plastid genomes, focusing on the green lineage that has given rise to three independent lineages with secondary plastids (euglenophytes, chlorarachniophytes, *Lepidodinium*). Our results show an overall increase in gene loss associated with secondary endosymbiosis, but this loss is tightly constrained by retention of genes essential for plastid function. The data show that secondary plastids have experienced temporary relaxation of purifying selection during secondary endosymbiosis. However, this process is tightly constrained as well, with selection relaxed only relative to the background in primary plastids, but purifying selection remaining strong in absolute terms even during the endosymbiosis events. Selection intensity rebounds to pre-endosymbiosis levels following endosymbiosis events, demonstrating the changes in selection efficiency during different phases of secondary plastid origin. Independent endosymbiosis events in the euglenophytes, chlorarachniophytes, and *Lepidodinium* differ in their degree of relaxation of selection, highlighting the different evolutionary contexts of these events. This study reveals the selection-drift interplay during secondary endosymbiosis, and evolutionary parallels during the process of organelle origination.

## Introduction

The endosymbiosis event leading to present-day chloroplasts is inferred to have taken place ∼1.5 billion years ago through the incorporation of a cyanobacterium by a heterotrophic host (Yoon, et al. 2004; Price, et al. 2012; Nowack and Weber 2018). This endosymbiosis event is referred to as primary endosymbiosis, with the plastids of the organisms descending from this event termed primary plastids (Keeling 2010; Archibald 2015). Three photosynthetic lineages emerged from this ancestor: the Chlorophyta (green algae), Rhodophyta (red algae) and Glaucocystophyta. Subsequently, several red and green algae have engaged in secondary endosymbiosis events, giving rise to more complex plastids. Secondary endosymbiosis differs in having a eukaryotic alga (carrying a primary plastid) as the photosynthetic partner being established as an organelle, and this process has spread photosynthesis to many unrelated branches of the eukaryotic tree of life (Keeling 2010). Despite the relevance of plastid endosymbiosis for eukaryotic evolution and algal diversity, the understanding of molecular evolution during the origination of these plastids is limited.

Endosymbionts often experience lowered levels of natural selection (Latorre and Manzano-Marín 2017; Wernegreen 2017), with the elevation of levels of stochastic genetic drift leading to an accumulation of slightly deleterious mutations, resulting in genome reduction and making them more susceptible to degradation (Moran 1996; Pettersson and Berg 2007; Moran, et al. 2008; Bennett and Moran 2015). Plastids have retained a highly reduced genome (ca. 100-200kb) characterised by accelerated rates of evolution and AT-biased nucleotide composition compared to free-living cyanobacteria (Green 2011; Bennett and Moran 2015). As is the case in many endosymbionts, plastid genomes have lost the majority of cyanobacterial genes, some having been transferred to the nucleus. Some of the gene losses are compensated by nucleus-encoded plastid-targeted genes that enable integration of plastids into the host cell biology. Plastid genomes have a highly conserved set of key genes encoding for core components involved in photosystem, ATP synthesis and protein translation(Allen 1993, 2017) that are under strong purifying selection (Smith 2015; Grisdale, et al. 2019). Several hypotheses suggest that the retention of genes in the plastid genome enhances the ability of organelles to efficiently respond to fluctuating conditions (Allen 1993, 2017; Johnston 2019). Strong purifying selection on the retained plastid genomes distinguishes them from most other endosymbiont genomes in early stages of endosymbiosis. While parallels can be expected between the evolutionary forces acting during establishment of plastid endosymbiosis (e.g. (Reyes-Prieto, et al. 2010; Lhee, et al. 2019) and other obligate endosymbiosis events based on the similarities in their overall genomic features, there has been very little work on characterising patterns of selection and drift in the origination of plastid organelles.

Secondary endosymbiosis differs fundamentally from primary because at the start of this process, the genomes of the primary plastid have already transitioned to a reduced state (Green 2011), with secondary green plastids having roughly similar gene content to primary green plastids (Suzuki, et al. 2016; Karnkowska, et al. 2018). Inouye and Okamoto (2005) postulate that secondary endosymbiosis of plastids involves several stages, beginning with permanent retention of the engulfed primary alga, followed by reduction of the endosymbiont genomes (primarily the nucleus) and ultimately fixed as an organelle through nuclear encoded plastid targeted genes. Recent studies have emphasized the possible role of the secondary host nucleus in facilitating the integration of the incoming green plastids in lineages that have hosted other plastids before (Ponce-Toledo, et al. 2018; Ponce-Toledo, et al. 2019). All these previous studies related to secondary endosymbiosis are focused on the endosymbiont’s nuclear genome reduction, but the molecular evolution of plastid genomes through the various stages of secondary endosymbiosis remains largely unexplored.

This study aims to characterise the molecular evolutionary processes acting on the origin of secondary plastids, using secondary plastids of green algal ancestry as a model system. These secondary plastids are found in three lineages, the chlorarachniophytes (a group of Rhizaria), the euglenophytes (a group of excavates) and the dinoflagellate genus *Lepidodinium* (Jackson *et al*. 2018). The existence of these three evolutionary events, distinctly independent from each other and with clearly identifiable host and plastid donor origins, makes green-type secondary plastid an excellent case study to investigate features common to secondary endosymbiosis events and those unique to individual events. Here, we use phylogenetic methods to examine the variation in selection on genes before, during, and after endosymbiosis, and to compare how this selection varies across genes and endosymbiosis events. We also quantify patterns and rates of gene loss across these events of secondary endosymbiosis. Our results are interpreted in the light of evolutionary processes that can contribute to variation in selection during secondary endosymbiosis.

## Results and Discussion

### Plastid genome features

Most plastid genomes, including those of secondary plastids, had small genomes (median 153kb), low GC proportion (0.34) and encoded an average of 80 annotated protein-coding genes. Plastid genomes of chlorarachniophytes (70kb genome, 60 CDS) and *Lepidodinium* (66kb, 62 CDS) are smaller with fewer CDS than those of euglenophytes plastid genomes (90kb, 64 CDS) (Table S1). Codon usage bias estimated using synonymous codon usage order showed that all green plastids studied had similar codon usage bias that appeared proportional to nucleotide composition (Figure S1). Among the secondary plastid lineages, chlorarachniophyte plastids had slightly lower GC content and higher codon usage bias than euglenophytes and *Lepidodinium*. However, codon usage bias for secondary plastids was within the range of that observed for primary plastids.

### Tightly constrained genome reduction

By grouping protein-coding genes into orthogroups and estimating gene loss with Dollo parsimony, it became apparent that plastid genomes experience an elevated level of genome reduction during secondary endosymbiosis events, but that they retain all key plastid genes encoding for core subunits related to photosynthesis, ATP and protein synthesis (Figure 1 and Figure 2). Reductive genome evolution highlights the similarities in molecular evolution between secondary plastid endosymbiosis and many examples of bacterial endosymbiosis in insects (McCutcheon and Moran 2012). Gene loss is particularly severe during primary endosymbiosis, with cyanobacteria-sized genomes (ca. 1,800-12,000 genes) reducing to the ca. 80-230 genes found in primary plastids (Gabr, et al. 2020). Gene loss from plastids during secondary endosymbiosis was small in comparison, with our estimates indicating that chlorarachniophytes lost 30 genes during secondary endosymbiosis followed by euglenophytes with 24 and *Lepidodinum* with 22 gene losses (Figure 1). Even though the endosymbiotic branches are among the top five branches losing the most genes, the difference compared to the background is not statistically significant (ANOVA and Tukey HSD tests), possibly due to the small sample size (n=3) of endosymbiotic branches available for analysis.

**Figure 1.**
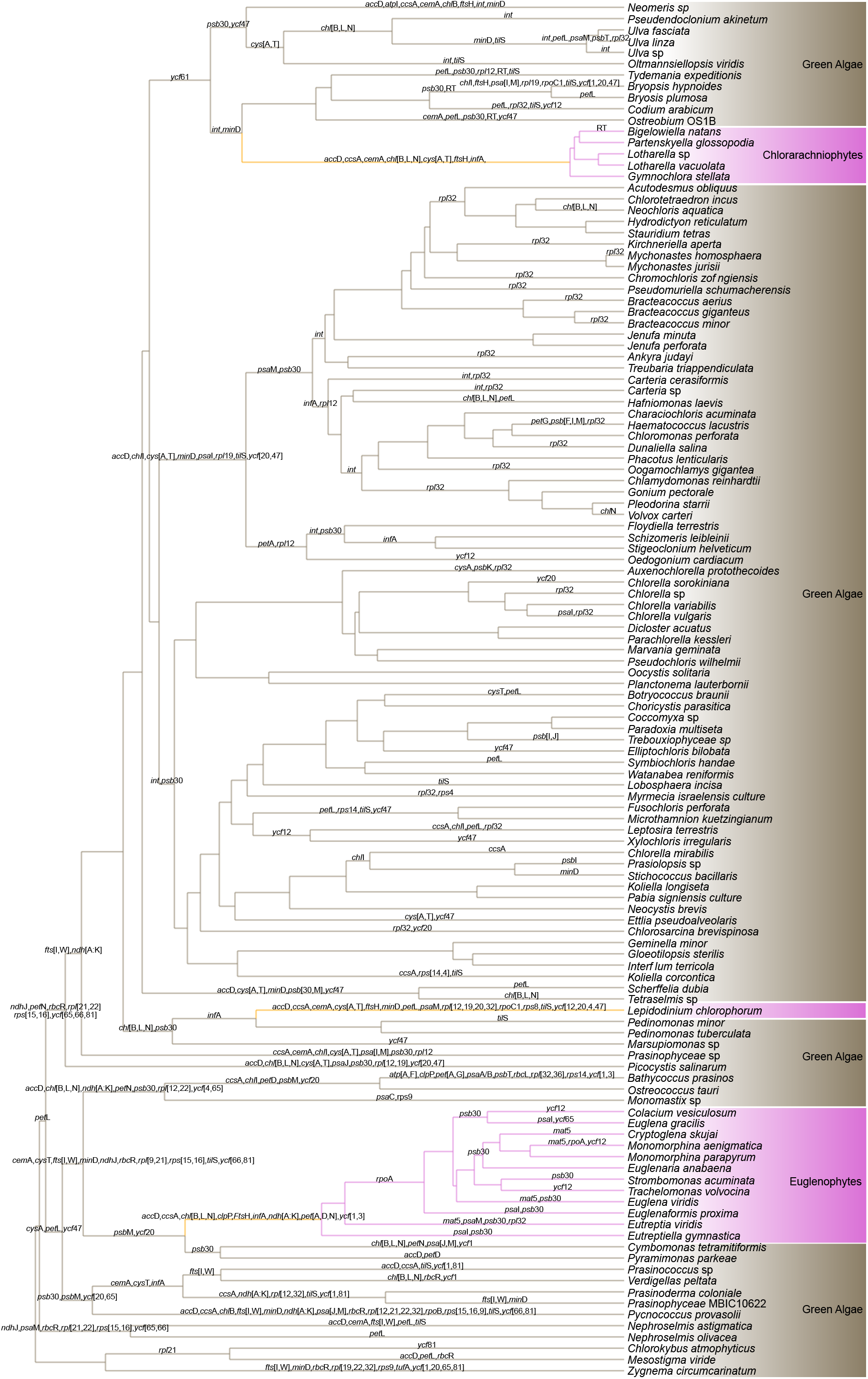
Evolution of green-type plastids across endosymbiosis events. The phylogeny is a chronogram indicating lineages as having primary plastids (i.e. green algae, in wheat brown), branches with secondary plastids (i.e. Chlorarachniophytes, *Lepidodinium*, Euglenophytes, in pink) and branches along which endosymbiosis happens (orange). Inferred gene losses are indicated along the branches.

**Figure 2.**
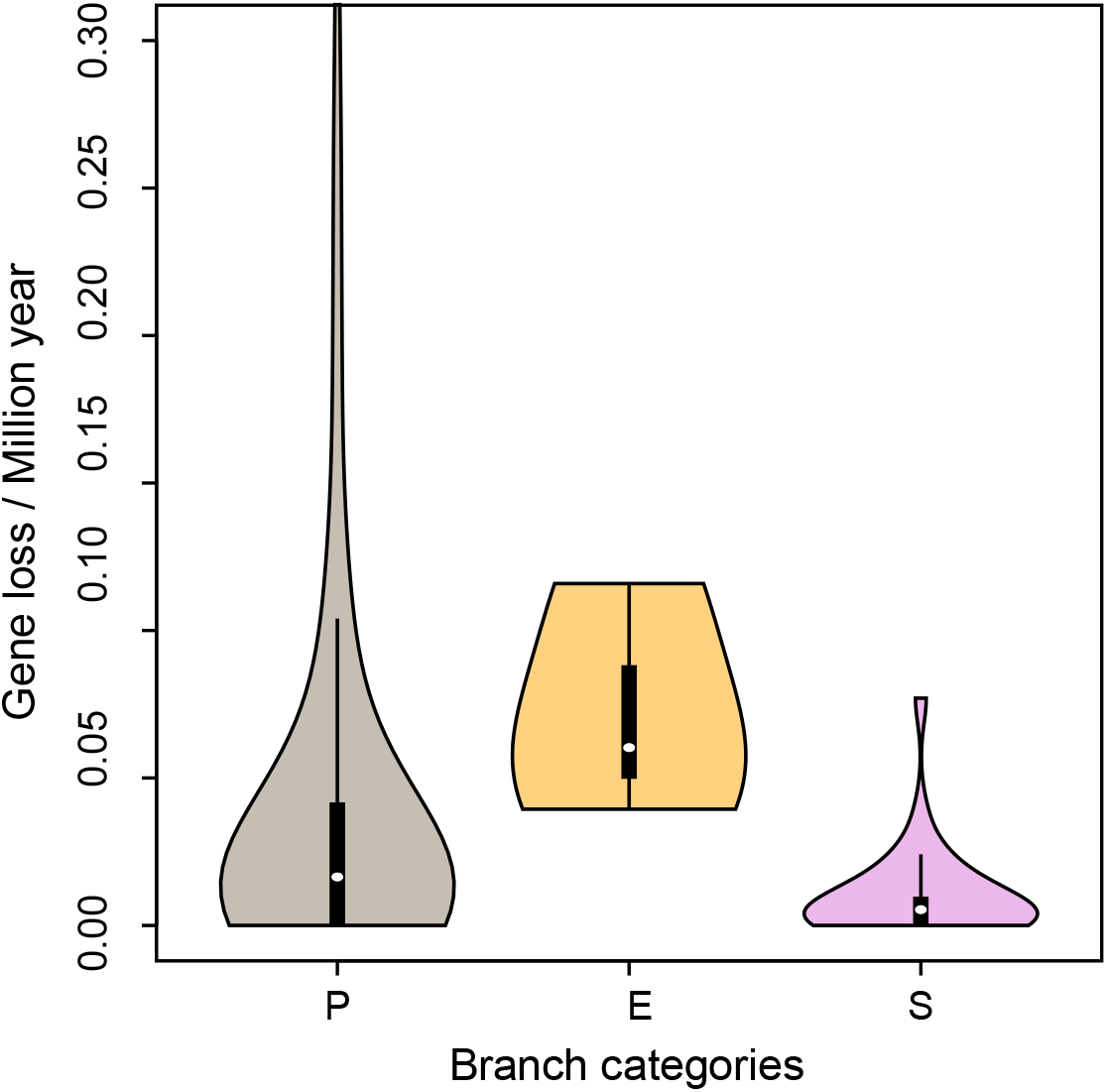
Inferred rates of gene loss in branches with primaryplastids(P),secondaryplastids(S) and branches along which endosymbiosis (E) takes place.

When viewed as the rate of gene loss per million years of evolution, the endosymbiosis branches had somewhat higher rates on average (Figure 2) but ranked lists showed that chlorarachniophytes and *Lepidodinium* were not among the branches losing genes fastest. So, despite most gene losses occurring on the endosymbiotic branches, the rates of loss per million years for these branches are not particularly high, suggesting that gene loss is a punctuated process occurring early in endosymbiosis (Moran and Mira 2001; Oakeson, et al. 2014). When correcting for the branch lengths of endosymbiotic branches, this punctuated effect is diluted to the point of not differing from background rates. Interestingly, the three independent endosymbiosis events showed similar numbers of gene losses (in the 22-30 range), adding to a list of similarities between secondary endosymbiosis events that also includes the convergent evolution of nucleomorph architecture seen in chlorarachniophytes and cryptophytes (Sarai, et al. 2020; Sibbald and Archibald 2020).

Our gene loss analysis showed that 17 genes were lost more than 10 times across the phylogeny, including *rpl*32 (ribosomal protein, 30 times), *psb*30 (photosystem II, 22), *til*S (tRNA Ile-Lysidine synthetase, 18), *pet*L (Cytochrome b6-f complex, 16) and *ycf*47 (14). Only *accD* (lipid acid synthesis), *ccsA* (mediates heme attachment to c-type cytochromes) and *ftsH* (cell division) were lost in all three endosymbiotic events. Some genes lost during one endosymbiotic event are also absent from other secondary plastids, but were lost before the endosymbiotic event. For instance, *ndh* [A:I,K](NAD(P)H oxidoreductase) was lost during euglenophyte endosymbiosis but it was also lost from the green algal lineages that gave rise to the chlorarachniophytes and *Lepidodinium*. Most of the genes lost during secondary endosymbiosis are likely to be compensated by nuclear homologs or through an alternative mechanism. For instance, the light-independent chlorophyll synthesis genes *chl*B, *chl*L and *chl*N that were lost during chlorarachniophyte and euglenophyte endosymbiosis and in many other primary plastids, including the ancestors of *Lepidodinium*, can be compensated by the light-dependent chlorophyll production pathway (Hunsperger, et al. 2015).The *chl*B, *chl*L and *chl*N genes have also been lost from some secondary plastids of cryptophyte algae (Fong and Archibald 2008).

Homologs of *rpl*12, *rpl*32, *rps*9 (small ribosomal proteins), *inf*A (translational initiation factor) and *fts*H were found in the nuclear genome of the chlorarachniophyte *Bigelowiella natans* (Curtis, et al. 2012), suggesting they may have been transferred from the plastid rather than lost entirely. Similarly, homologs of *pet*A, *pet*N, *ycf*3, *clp*P (Clp protease proteolytic subunit) and *fts*H were recovered in the transcriptomes of the euglenophytes *Euglena gracilis* and *Eutreptiella* (Hrdá, et al. 2012). Aside from the genes mentioned above, all other genes lost during secondary endosymbiosis including genes with a function in photosynthesis (like *psb*30, *psb*M, *psa*I) were not detected in the nuclear genomes of *B. natans* and *E. gracilis* and may represent genuine gene losses, but some caution is warranted as most of these proteins are small and may be missed in genome-wide blast searches.

Several of the genes predicted to be lost during secondary endosymbiosis (*acc*D, *inf*A, *ndh, ycf*1, *ycf*3 and *ycf*4) were lost from plastid genomes in other lineages too, and compensatory nuclear-encoded genes have been identified (Boudreau, et al. 1997; Millen, et al. 2001; Martín and Sabater 2010; Huerlimann and Heimann 2012).

Losses of genes whose functions can be compensated are likely to have little impact on plastid function. Loss of similar genes in parallel in different parts of the tree suggests they may experience reduced selective constraints compared to key photosynthesis genes, and in periods with increased drift, such genes may be more likely to be lost than genes under stronger selection. Recent work shows that genes encoding central subunits of the electron transport chain are more likely to retained in the organelle (Johnston and Williams 2016). In line with this, we see that across the green algal phylogeny, 48 genes including the core components of photosynthesis and protein synthesis remained highly conserved (never lost or lost once).

The role of selection in retaining genes has also been demonstrated in the chromatophore genomes of *Paulinella*, a model species for the study of primary endosymbiosis (Reyes-Prieto, et al. 2010; Valadez-Cano, et al. 2017; Lhee, et al. 2019). Overall, our results suggest that genome reduction appears to be elevated during secondary endosymbiosis but is a tightly constrained process with strong selection to retain genes with key functions. Of course, the lineages with secondary endosymbionts that we study here are all photosynthetic, implying that by the design of our study, we introduced a bias towards endosymbiosis events that would have maintained all genes with an essential function in photosynthesis. It is perfectly conceivable that other outcomes are possible in endosymbiosis events that do involve loss of photosynthetic function, but we are not aware of any instances where a cyanobacteria or eukaryotic alga has been retained as an endosymbiont for functions other than photosynthesis, besides secondarily non-photosynthetic groups such as apicomplexans.

### Selection dynamics through endosymbiosis

For our analysis of selection dynamics through endosymbiosis, the phylogeny was divided into three sets of branches representing primary plastids (P), secondary plastids (S) and endosymbiosis branches (E). Selection intensity during secondary endosymbiosis was quantified using a Hyphy RELAX model that contrasts the selection on the endosymbiosis branches relative to all other branches. The relative selection intensity parameter (k-value) of the fitted model showed that the distribution of k-values across genes is well below 1 (median 0.43), a clear signature of relaxation of selection in the endosymbiotic branches (E) compared to all other branches (P+S) (Figure 3, and see Tables 1, S2). Of the 34 genes in the analysis, 26 showed statistically supported relaxation. Two outlier genes (*psb*D and *psb*E) showed slight intensification of selection (k>1) for this model, but without significant statistical support. The same model (denoted E × P+S) applied to a concatenated alignment of all plastid genes (Table S3) returned results in line with the findings for individual genes, with relative selection intensity parameter (k) value of 0.55. The E × P+S model is a significantly better fit to the concatenated sequences than the null model (p<0.0001 and likelihood ratio= 557.65), implying a significant decrease in evolutionary selection (relaxation) during endosymbiosis.

**Table 1:** The number of green algal plastid genes showing numbers of genes showing relaxation and intensification for each model.

| Model | Relaxation<br>($k < 1$) | Significant<br>Relaxation | Intensification<br>($k > 1$) | Significant<br>Intensification | Neither<br>$k = 1$) |
| --- | --- | --- | --- | --- | --- |
| **E × P+S** | 32 | 26 | 2 | - | - |
| **S × P(E)** | 19 | 13 | 12 | 9 | 3 |

**Figure 3.**
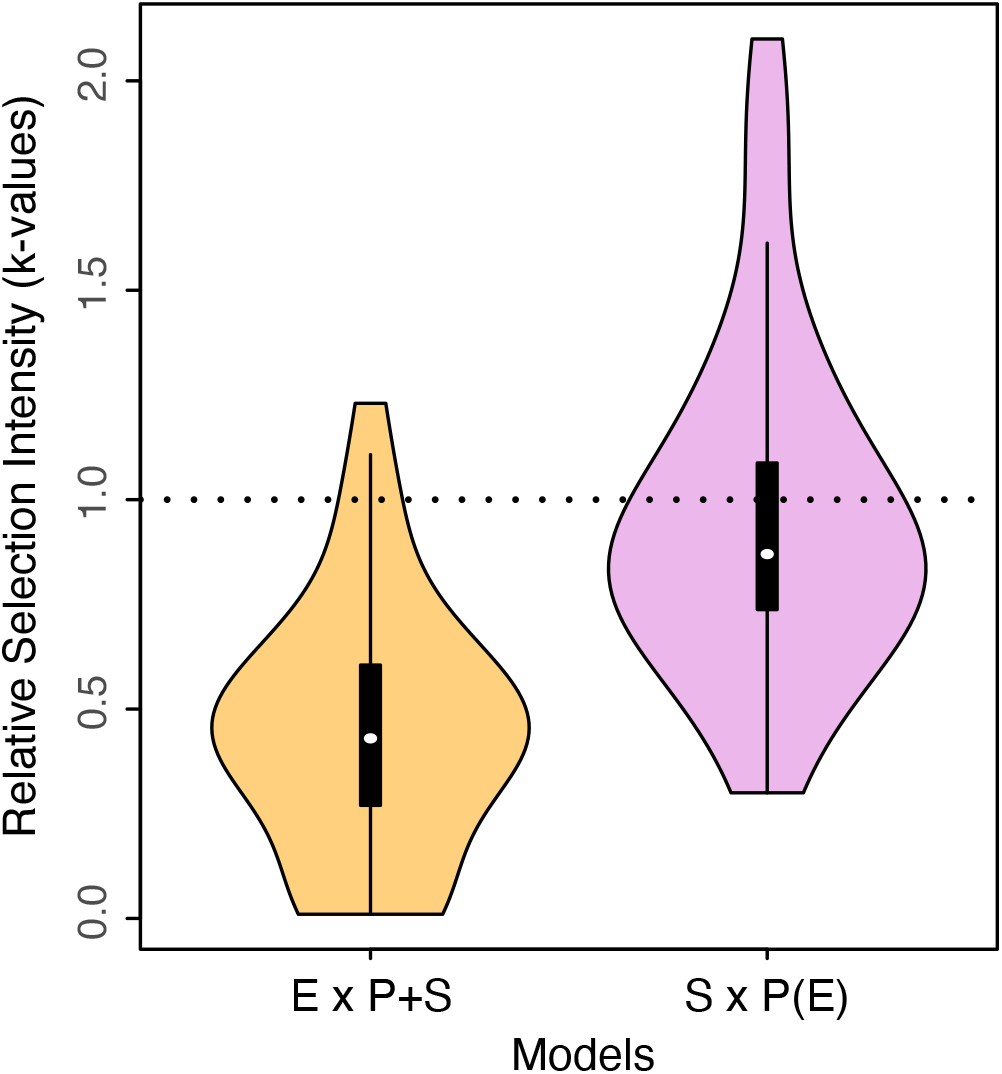
Distribution of the relative selection intensity parameter (k) of the HyPhy RELAX model for (i) Endosymbiosis (test) vs. Primary and Secondary branches (reference) [E × P+S model],(ii) Secondary (test) vs. Primary branches (reference), excluding endosymbiosis branches [S × P(E) model]. Selection intensity is relaxed in the test branches when k<1 or intensified when k>1. These plots show that endosymbiosis branches have relaxed selection compared to primary and secondary branches, and that selection on secondary branches is similar to that of primary branches, indicating that the relaxation of selection during endosymbiosis is temporary.

While the signature of relaxation is clear, this does not imply that molecular evolution is neutral in endosymbiotic branches. The model categorised 82.12% of sites as being under purifying selection, with ω (ratio of non-synonymous(dN) to synonymous(dS) substitutions) value of 0.06 in the endosymbiotic branches indicating that most sites remain under purifying selection even during endosymbiosis events. BUSTEC analyses provided additional statistical support for purifying selection along endosymbiosis branches, with all genes having lower AIC scores for the unconstrained model with purifying selection than for the model constrained to exclude purifying selection (Table S4).

Selection analysis based on the E × P+S model and the BUSTEC results helps to characterise the molecular evolutionary process during secondary plastid endosymbiosis. Studies of insect endosymbionts suggest that relaxation of purifying selection during endosymbiosis establishment in obligate endosymbionts of insects can be due to two processes: a population bottleneck and decrease in functional constraints on proteins (Moran 1996; Wernegreen 2004, 2015). In the case of secondary plastid endosymbiosis, it seems unlikely to have much relaxation on functional constraints, in line with the observations of purifying selection and tight constraints on gene loss. Also, the relaxation is observed on nearly all retained genes, further shifting the balance of evidence towards population size effects on plastid genomes evolution during endosymbiosis. The near-neutral theory predicts that in small populations, the fate of near-neutral mutations depends on the balance between selection and the stochastic effect of drift (Ohta 1972, 1992). During bottlenecks, one can expect strongly deleterious mutations to continue being eliminated, while slightly deleterious mutations will have higher chances of being fixed in the population by stochastic drift than being eliminated by selection (Woolfit and Bromham 2003). In the chloroplast genes studied here, one would expect this process to result in more non-synonymous substitutions in the endosymbiotic branches, in line with the reduced selection efficiency we observe.

Relative selection analysis using a different model comparing secondary plastids to primary plastids (denoted S × P(E)) suggests that relaxation of selection during endosymbiosis is temporary, indicated by distribution of k-values that encompasses 1 (median 0.87) and similar numbers of genes that were relaxed (13), intensified (9) or inconclusive (9) in secondary branches. The analysis on concatenated sequences showed similar results (median k = 0.96) and was not preferred over the null model, providing a clear indication that following the relaxation during endosymbiosis, the purifying selection regime on plastid genes returns to values similar to those before endosymbiosis.

Comparative studies of the genomes of endosymbionts at different stages of integration have shown that genome stability increases with the age of the endosymbiont and suggested this may be due to selection (Allen, et al. 2009; Martínez-Cano, et al. 2015). Our findings agree with these observations, and our model system has the added advantage of the endosymbiont becoming a stable organelle, fully integrated and co-diversifying with the host following endosymbiosis, which was not the case in the previously studied endosymbiont models. This allowed us to disentangle the molecular dynamics along the endosymbiosis branch from that of a stable integrated secondary plastid, showing that the purifying selection regime rebounds to near pre-endosymbiosis levels once the organelle is established.

Our results suggest a general model for the molecular dynamics of secondary plastid endosymbiosis (Figure 4). It is likely that a very small fraction of the actual population of the engulfed primary alga is involved in secondary endosymbiosis, creating a drastic population size bottleneck. This decrease in effective population size would then allow higher levels of drift to fix slightly deleterious mutations, explaining the long branches in the phylogeny of green plastids where secondary endosymbiosis events take place (Jackson et al. 2018).

**Figure 4:**
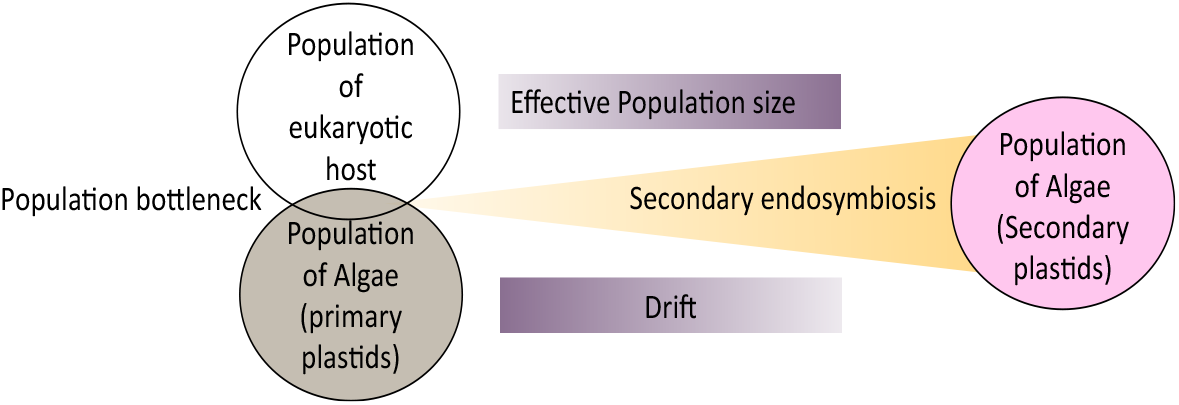
A general model for molecular dynamics during secondary green-type plastid endosymbiosis. The model illustrates the population bottleneck due to involvement of very small fraction of the actual population of the engulfed primary alga in secondary endosymbiosis. As the endosymbiont-host relation ages the effective population size increases that counteracts the impact of stochastic drift leading establishment of secondary plastids after endosymbiosis.

Maintenance of the plastid genome during secondary endosymbiosis depends largely on nuclear-encoded DNA replication and repair proteins (Smith and Keeling 2015). During secondary endosymbiosis, nuclear-encoded proteins are often transferred from the algal nucleus to the new host nucleus, with the product directed to the new plastid. This might contribute to a period of reduced fidelity of plastid DNA replication during secondary endosymbiosis, which might in some cases lead to failure of the secondary plastid endosymbiosis. As the endosymbiont-host relationship ages, the drift acting on plastid genomes could eventually decrease, with higher effective population size and level of integration of plastid and host nucleus. This is reflected in the increased levels of selection on secondary plastids following endosymbiosis, emphasising the important interplay between drift and selection during secondary endosymbiosis and their resulting impact on secondary plastid genomes.

### Three independent events

Our analyses comparing selection regimes of the three endosymbiosis events to the background individually showed distinctive scenarios. *Lepidodinium* showed the strongest relaxation (k = 0.3) followed by chlorarachniophytes (k = 0.45), indicating evidence of strongly relaxed selection during these two endosymbiotic events. However, euglenophytes showed a k-value of 0.86, indicating a much lower level of relaxation during this endosymbiosis event.

Tightly constrained genome reduction along with evident purifying selection across all three green algal secondary endosymbioses emphasises the evolutionary parallels among these independent events, but also clearly distinguishes the origin of secondary green plastids from other recently established obligate endosymbionts. Differences in degrees of relaxation and gene losses during these three secondary endosymbiosis events highlight the different evolutionary pressures associated with them. Nutritional requirements and level of mixotrophy can account for different evolutionary pressures during plastid endosymbiosis.

Because our analyses support increased drift, the differing relaxation intensity between the events implies that there may be differences in the extent of population bottlenecks underlying these events. Among the three events, euglenophytes are noticeable as they had the least relaxation of selection. The endosymbiosis branch leading to the euglenophytes in the phylogeny is shorter than the other two endosymbiosis branches, suggesting that plastid genes have evolved through this secondary endosymbiosis with fewer substitutions (Jackson et al. 2018, Figure 1). This might be due to a less intense population bottleneck with more efficient integration of the plastid during euglenophyte endosymbiosis compared with the other two green secondary endosymbioses. Absence of homologs of plastid origin for protein import components and plastid division in euglenophytes has led to speculation that integration of their plastids involved a novel/simplified process including proteins of host origin (Zahonova, et al. 2018; Novák Vanclová, et al. 2020), which could have facilitated more efficient integration of their plastid genomes, allowing faster recovery from bottleneck. This may have enabled euglenophyte plastids to integrate with less relaxation of selection.

## Material and methods

### Dataset

We compiled a dataset of 122 green plastid genomes spanning the primary plastids of green algae (Chlorophyta, 104 genomes) and the secondary plastids of Euglenophyta (12 genomes), Chlorarachniophyta (5 genomes) and the dinoflagellate genus *Lepidodinium* (1 genome). A reference phylogeny (Figure 1) was obtained from a previous study (Jackson, et al. 2018). Our dataset includes close extant relatives of ancestral green algae that were involved in the secondary endosymbiosis events, making this green plastid dataset suited to examining the molecular evolutionary dynamics associated with secondary endosymbiosis, and investigating differences and similarities among the three independent cases of secondary green plastid origination. Basic features of the plastid genomes such as number of coding sequences (CDS) and genome size were recorded and GC content of CDS and codon usage bias were calculated using the CodonO (Wan, et al. 2007) function from the cubfits v.0.1-3 (Chen 2014) package in R v.3.5.1 (R core Team, 2018).

### Analysis of gene loss

To investigate evolutionary patterns of gene loss, protein-coding genes were grouped into orthogoups using OrthoFinder version 1.4.0 with standard parameters (Emms and Kelly 2015). Of the 203 orthogroups (OGs) that were present across multiple species, 116 OGs corresponded to named genes with known function conserved across most plastids, while the remaining OGs (mostly hypothetical genes of unknown function) were not examined further. A presence/absence matrix of the 116 orthogroups corresponding to named genes was constructed. Using this matrix and the reference phylogeny from Jackson et al. (2018), gene gain and loss along the phylogeny was estimated using PHYLIP version 3.695 (Felsenstein 2005), with the Dollo parsimony method and printing the states at all nodes of the tree. Gene loss and gain along each branch was extracted from the PHYLIP output using OrthoMCL Tools (DOI 10.5281/zenodo.51349). The rate of gene loss and gain per million years was calculated for each branch using the evolutionary time from the chronogram presented by Jackson et al (2018). The estimated numbers of genes lost (and rates of gene loss) were ranked from largest to smallest to see if endosymbiotic branches had greater values compared to the background, and evaluated formally using ANOVA and Tukey HSD tests in the stats v3.6.2 package of R core Team (2013). To investigate if the genes lost during the secondary endosymbiosis may have been transferred to host nuclear genomes, we performed local tBLASTn searches using orthologous genes as query against the published nuclear genomes of *Bigelowiella natans* (Curtis, 2012) and *Euglena gracilis* (https://www.ncbi.nlm.nih.gov/assembly/GCA_900893395.1) (e-value cut-off = 1e-05).

### Selection intensity analysis

To study the variation in selection intensity in the protein-coding genes of secondary and primary green plastids, we used the hypothesis-testing framework RELAX (Wertheim, et al. 2014) from the HyPhy software package version 2.3.14 (Kosakovsky Pond, et al. 2005; Delport, et al. 2010). This framework requires a predefined tree with subsets of test and reference branches specified. The subset of branches that are not set as test or reference remain unclassified. RELAX applies a branch-site model to estimate the strength of natural selection based on the ratio of non-synonymous to synonymous substitutions (omega, ω) for three different ω categories (ω_1_ < ω_2_ ≤ 1< ω_3_) in the test and reference subsets. ω < 1 represents sites under purifying selection, ω > 1 represents sites under positive selection and ω = 1 represents sites under neutral evolution. The relative selection intensity parameter (k) reflects intensification or relaxation of selection based on the relative proximity of ω values to 1 (neutral evolution). If ω values of test branches are closer to 1 than reference branches, then selection is relaxed (k<1) and in the opposite scenario, selection has intensified (k>1). The null model assumes identical ω values (k=1) between test and reference branches. The alternative model fits different sets of ω values for test and reference, and thus k differs from 1, allowing a formal test of relaxed (k<1) or intensified (k>1) selection. The likelihood ratio test(LR) performed with p-value < 0.05 by comparing the null and alternate model quantifies statistical confidence for the obtained k value.

### Models for Selection Analysis

We used the HyPhy-RELAX method to study molecular evolution through the process of endosymbiosis by designing different evolutionary models that allowed us to study aspects of selection intensity before, during and after the endosymbiosis process. For the selection analyses we included only genes that were present in all of the lineages with secondary plastids (34 orthologous genes). The phylogenetic tree of algal green plastid genomes from Jackson et al. (2018) was used as the predefined tree on which test and reference branches were marked. In the phylogeny (Figure 1), branches leading to and connecting the species containing primary plastids (i.e. the green algae) were indicated as primary branches (P), and denote the state before secondary endosymbiosis. Secondary branches (S) are the branches leading to and connecting the species containing secondary plastids, and denote the state after secondary endosymbiosis. The endosymbiotic branches (E) indicate branches connecting the backbone of green algal lineages to the lineages with secondary green plastids, in other words the branches along which secondary endosymbiosis took place (orange-coloured branches in Fig. 1). The *Lepidodinium* lineage includes only one plastid genome so we consider this branch as the endosymbiotic branch for this case.

Our first model, denoted “E × P+S”, has endosymbiotic (E) branches as the test set and all non-endosymbiotic branches (P+S) as the reference set. This model allows us to compare the selection intensity during endosymbiosis relative to before and after endosymbiosis. Our second model, denoted “S × P(E)”, allowed us to evaluate differences in selection intensity between secondary (S) and primary (P) plastids, excluding the endosymbiont branches (E).

To study differences between individual endosymbiosis events, we fitted E x P+S models, but specifying only a single endosymbiotic branch as test (excluding all other endosymbiotic branches) and all non-endosymbiotic branches (P+S) as the reference set.

### Purifying selection analysis

Because functional plastid genes are expected to experience purifying selection, we also carried out an analysis to identify and quantify levels of purifying selection. The BUSTEC method implemented in HyPhy tests for alignment-wide evidence of conservation by fitting a random effects branch-site model to the entire phylogeny or a subset of tree branches (Murrell, et al. 2015). The null model constrains ω values to greater than or equal to 1, excluding the possibility of purifying selection. The unconstrained model allowing ω values greater than and less than 1 serves as the alternate model. With endosymbiotic branches as the test branches, we used BUSTEC to fit the alternative unconstrained and null constrained models to these branches to quantify evidence for purifying selection during endosymbiosis.

## Funding

This work was supported by Australian Research Council Discovery Project to HV, CXC & DB (DP150100705).

## Acknowledgements

We thank the people who worked on chloroplast genomics and delivered the data needed for this study. We acknowledge the Traditional Owners of the land on which we work, and pay our respects to their Elders, past, present and emerging. We are thankful to Sergei L. Kosakovsky Pond for his recommendations on BUSTEC.

## Conflicts of Interest

The authors declare no conflicts of interests.

## Supplementary Materials

https://doi.org/10.26188/14616999

## Notes

### Competing Interest Statement

The authors have declared no competing interest.

